# Longitudinal Deep Multi-Omics Profiling in a *CLN3^Δex7/8^* Minipig Model Reveals Novel Biomarker Signatures for Batten Disease

**DOI:** 10.1101/2023.09.20.558629

**Authors:** Mitchell J Rechtzigel, Brittany Lee, Christine Neville, Ting Huang, Alex Rosa Campos, Khatereh Motamedchaboki, Daniel Hornburg, Tyler B Johnson, Vicki J Swier, Jill M Weimer, Jon J Brudvig

## Abstract

Development of therapies for CLN3 Batten disease, a rare pediatric lysosomal storage disorder, has been hindered by the lack of etiological insights and translatable biomarkers to clinics. Here, we used a deep multi-omics approach to discover new biomarkers using longitudinal serum samples from a porcine model of CLN3 disease. Comprehensive metabolomics was combined with a nanoparticle-based LC-MS-based proteomic profiling coupled with TMTpro 18-plex to generate quantitative data on 769 metabolites and 2,634 proteins, collectively the most exhaustive multi-omics profile conducted on serum from a porcine model, which was previously impossible due a to lack of efficient deep serum proteome profiling technologies compatible with model organisms. The presymptomatic disease state was characterized by elevations in glycerophosphodiester species and lysosomal proteases, while later timepoints were enriched with species involved in immune cell activation and sphingolipid metabolism. Cathepsin S, Cathepsin B, glycerophosphoinositol, and glycerophosphoethanolamine captured a large portion of the genotype-correlated variation between healthy and diseased animals, suggesting that an index score based on these analytes could have great utility in the clinic.

## Introduction

Batten Disease (also known as neuronal ceroid lipofuscinoses (NCLs)), are a group of neurodegenerative lysosomal storage disorders that result from pathogenic variants in one of 13 ceroid lipofuscinosis neuronal (CLN) genes. Collectively, Batten disease affects approximately 1 in 100,000 individuals worldwide, making it the most common pediatric neurodegenerative disorder^1^. The most common form of Batten Disease, CLN3 disease, is a rare and fatal autosomal recessive disorder caused by mutations in *CLN3*. Individuals with CLN3 disease typically experience vision loss in early childhood, followed by seizures, motor and cognitive decline, and premature death by the third decade of life^2, 3^. Pathologically, CLN3 dysfunction cascades from to the accumulation of lysosomal storage material, microglia and astrocyte activation, to neuronal dysfunction and death^4^ .

Despite many years of research, the molecular function of CLN3 and many of the other NCL proteins has yet to be fully elucidated. Recent advancements have begun to outline the biological processes affected by CLN3 dysfunction, but the field still lacks robust biomarker signatures that comprehensively reflect the disease state^5, 6^. With the growing list of CLN3-specific therapies entering clinical trials, there is a substantial need for non-invasive biomarkers that can track disease progression and therapeutic efficacy^2, 7^. We recently identified a group of glycerophosphodiesters as promising blood-based biomarker candidates for CLN3^8^. Shortly thereafter, a comprehensive series of biochemical experiments demonstrated that CLN3 is required for the clearance of glycerophosphodiesters from lysosomes^6^. Recent work also demonstrated that closely related phosphoinosides mediate lysosomal repair suggesting that disrupted glycerophosphodiester metabolism or transport could underlie the severe lysosomal dysfunction that characterizes cellular disease pathology^9^. Although elevation of these glycerophosphodiester species closely corresponds with the absence of functional CLN3, this phenotype does not correlate with other progressive aspects of disease progression such as neuroinflammation and neurodegeneration. In contrast, markers of neurodegeneration such as neurofilament light (NFL) show highly variable elevations in CLN3 disease and thus have questionable utility as diagnostic and prognostic biomarkers^10^ Overall, it is unlikely that any single biomarker will be adequate to evaluate the complex environment of disease status and progression in any individual patient. A combined biomarker “score” that integrates diverse sets of markers reflecting different facets of disease etiology and pathology could provide greater precision in tracking disease progression, accelerating therapeutic development.

We sought to uncover a more diverse set of CLN3-related biomarkers and to gain insights into the molecular function of CLN3 using an untargeted metabolomics and a novel deep multi-nanoparticle-based proteomics in a Yucatan Minipig model of CLN3 Batten disease harboring the most common patient mutation: a ∼1kb deletion in exons seven and eight^11, 12^(Fig. 1). We analyzed blood serum samples from the model animals at three stages of disease progression to capture temporal changes and disease-specific patterns in the proteome and metabolome at both the pathway and individual molecule levels. In contrast to tissue biopsies, sampling blood can serve as a minimally invasive procedure for monitoring disease progression, enabling comprehensive proteomic research and in-life monitoring of biomarker status. However, historically the extreme dynamic range of blood protein concentrations has required a trade-off between depth of unbiased proteome coverage and number of samples analyzed, in particular for model organisms that lack abundant protein depletion solutions and that are not compatible with targeted strategies based on aptamers and antibodies designed for human proteomes^13^. Here, we utilized a nanoparticle-based protein sampling technology, the Proteograph^TM^ Product Suite (Seer, Inc.), enabling deep and scalable proteomic profiling, independent of disease model species^11, 14, 15^. Our in-depth profiling of thousands of proteins with more than 10,000 peptides, integrated with in-depth metabolome data, uncovered novel signatures and biomarker candidates for CLN3 disease and provided insights into perturbed pathways in the CLN3 disease state.

**Figure 1:**
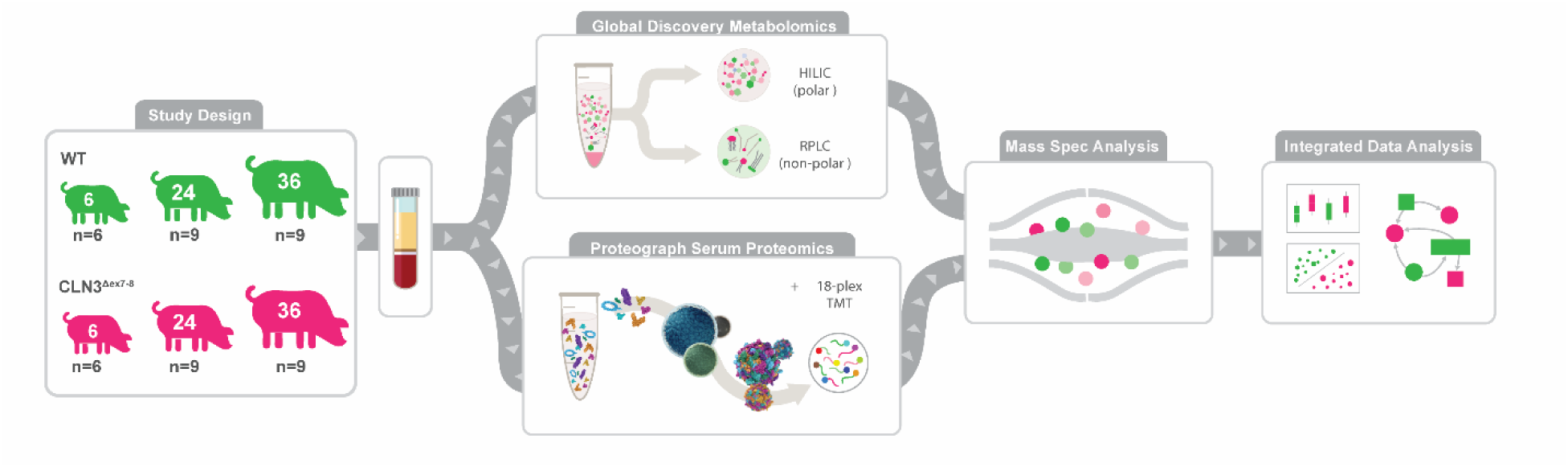
Study Design and Multiomics Analysis Workflow. Serum samples were taken from control and *CLN3*^Δex7–8^ Yucatan miniature pigs at 6-, 24-, and 36-months. One aliquot of serum was analyzed with the Global Discovery Panel: a comprehensive mass spectrometry analysis including both metabolites and lipids. Another aliquot of serum was used to perform deep proteomics using Seer’s nanoparticle technology and labeling with Tandem Mass Tag, high throughput mass spectrometry method. After mass spectrometry analysis of each omic data, the data was processed and integrated for statistical, multivariate analyses.

## Methods

### Animal Ethics Statement

Wild type (WT) and transgenic *CLN3^Δex7–8^* Yucatan miniature pigs were housed and maintained at Precigen Exemplar under an approved Institutional Animal Care and Use Committee (IACUC) protocol.

### CLN3^Δex7–8^ Mini Pig Generation

Transgenic *CLN3^Δex^*^7–8^ mini pigs were generated as previously described^8^. In brief, wild type fetal Yucatan mini pig fibroblasts were transduced with a recombinant AAV1 containing a *CLN3^Δex^*^7–8^-Neo targeting vector that covered exon six to intron nine while excluding a ∼1-kilobase region spanning exon seven and eight, mirroring the most common CLN3 disease mutation. Following antibiotic selection, PCR-positive clones underwent treatment with a recombinant AAV1 containing a Cre recombinase expression cassette to excise the integrated selection cassette. Southern blot and sequencing were employed to screen recombinant clones and identify those harboring on-target integrations. Nuclear transfer and embryo transfer were conducted at Precigen Exemplar Genetics (Germantown, Maryland, USA). Pregnancy of recipient animals was verified via abdominal ultrasound at day 21 and throughout gestation. Study animals were bred *CLN3*^Δex7–8^ heterozygote to *CLN3*^Δex7–8^ heterozygote, and genotype confirmed by PCR.

### Pig Biofluid Collection

As previously described, pigs were anesthetized with xylazine (TKX) and isoflurane (1-2%)^8^. Briefly, a 16 G needle attached to a 20-cc syringe was inserted into the right ventricle of the heart, and approximately 20mL of blood was drawn. A Saf-T Holder™ transfer device was used to expel collected blood into two 10mL Monojet™ blood collection tubes. Blood samples were placed at room temperature for 30 minutes and allowed to clot, after which they were centrifuged at 3100 x g for 10 minutes at room temperature. Serum was then collected into 2 mL polypropylene screw-top tubes and stored at −80 °C. Samples were collected at 6-months ± 7 weeks, 24-months ±10 weeks, 36-months ± 6 weeks, and 48-months ± 12 weeks in both *CLN3*^Δex7–8^ (n = 6; 9; 9; 3; respectively) and control (n = 6; 9; 9; 3; respectively) mini pigs.

### Untargeted Metabolomics Discovery

#### Metabolomic Sample Preparation

Metabolomic analyses were performed by Metabolon (Morrisville, North Carolina, USA) as previously described^8^. Briefly, samples were shipped on dry ice overnight to Metabolon. Samples were inventoried and promptly stored at −80°C until analyzed. Automated sample preparation was conducted using the MicroLab STAR® system (Hamilton Company) utilizing several recovery QC standards preceding the extraction process. Proteins were precipitated with methanol in a Genogrinder 2000 (Glen Mills) followed by centrifugation. Samples were divided into five fractions (two aliquots for replicate analyses with Reversed Phase Ultrahigh Performance Liquid Chromatography coupled to Mass Spectrometry (RP)/UPLC-MS/MS using positive ion mode electrospray ionization (ESI), one aliquot for RP/UPLC-MS/MS, with negative ion mode ESI, one aliquot for HILIC/UPLC-MS/MS in negative ion mode ESI, and one sample aliquot was reserved for backup). The organic solvent was removed via TurboVap® (Zymark). Samples were stored overnight in liquid nitrogen prior to LC-MS/MS analysis.

#### Quality Assurance for Metabolomic Analysis

To monitor LC-MS instrument performance and chromatographic alignment, internal controls were included with experimental samples for each run. As previously described, these included pooled matrix controls of well-characterized human serum, process blanks consisting of extracted water samples, and a cocktail of QC standards that were selected not to interfere with the measurement of endogenous compounds^8, 16, 17^. Median relative standard deviation (RSD) was calculated for the QC standards to monitor instrument variability from run to run. Median RSD was calculated for all endogenous metabolites (i.e., non-instrument standards) present in 100% of the pooled matrix samples to account for overall process variability. QC samples were spaced evenly among the injections with experimental samples randomized across the platform run.

#### Ultrahigh Performance Liquid Chromatography-Tandem Mass Spectroscopy Analysis for Discovery Metabolomics

As previously described, ultra-performance liquid chromatography coupled to tandem mass spectrometry (UPLC-MS/MS) analysis was conducted on a Waters^TM^ ACQUITY^TM^ UPLC and a Thermo Fisher Scientific^TM^ Q Exactive^TM^ Orbitrap^TM^ high resolution and accurate mass spectrometer interfaced with a heated electrospray ionization (HESI-II) source and Orbitrap mass analyzer operated at 35,000 mass resolution^8^. Samples were dried and reconstituted in method-compatible buffers for each of the LC-MS/MS acquisitions, which contained internal standards at fixed concentrations to control for injection and chromatographic run variations. For LC-MS/MS runs conducted in acidic positive ion conditions, chromatographically optimized for more hydrophilic compounds, the extracts were applied to a C18 column (Waters UPLC BEH C18-2.1×100 mm, 1.7 µm) followed by isocratic elution using water and methanol, containing 0.05% perfluoropentanoic acid (PFPA) and 0.1% formic acid (FA). Additionally, a separate aliquot was analyzed using acidic positive ion conditions; however, the extract was gradient eluted from the same C18 column using methanol, acetonitrile, water, 0.05% PFPA and 0.01% FA and was operated at an overall higher organic content. For samples analyzed using basic negative ion optimized conditions, a separate dedicated C18 column was utilized, and extracts were gradient eluted using methanol and water, however with 6.5mM ammonium bicarbonate at pH 8. The fourth aliquot was analyzed via negative ionization following elution from a hydrophilic interaction liquid chromatography (HILIC) column (Waters UPLC BEH Amide 2.1×150 mm, 1.7 µm) using a gradient consisting of water and acetonitrile with 10mM ammonium formate, pH 10.8. The MS analysis alternated between MS and data-dependent MS^n^ scans using dynamic exclusion. The scan range covered 70-1000 m/z, varying slightly across methods. Raw data files were extracted and analyzed as described in the data analysis section.

#### Metabolomic Data Extraction and Compound Identification and Quantification

Metabolon’s hardware and software systems were used for raw data extraction, peak identification, and quality control processing. Compounds were identified by comparing their retention time/index (RI), mass to charge ratio (m/z), and tandem MS/MS spectral data matched to library entries of purified standards or recurrent unknown entities. Metabolites were quantified using area-under-the-curve.

#### Metabolomics Data Processing

To estimate relative abundances, metabolite chromatographic peak area data was log_2_-transformed, then values for each individual metabolite were scaled across samples to a mean of zero and unit variance. No batch normalization was necessary as all metabolites were detected in one run. Metabolites missing in more than 50% of the samples were removed from consideration, leaving 769 metabolites in the dataset. We present an in-depth analysis of data used in our previous work, however comparisons and analyses made here are novel^8^.

### Targeted Protein Analysis

Neurology 4-PlexA targeted proteomic analysis was performed in singlet at a 4:1 sample dilution at the Simoa® Accelerator Laboratory (Billerica, Massachusetts, USA). This targeted proteomics panel included data for four proteins (GFAP, NFL, TAU, and UCHL1) and was processed in the same manner as the metabolomics data (Supplementary Fig. 1).

### Deep Proteomics Analysis

#### Automated Blood Serum Sample Processing with Proteograph^TM^ Workflow

Samples were processed by SP100 automation instrument with Proteograph^TM^ Assay Kt included in Proteograph Product Suite (Seer, Inc.) using five distinctly functionalized nanoparticles (NPs). In the fully automated workflow, 250µL of serum were equally aliquoted into 5 tubes where 40 µL of serum from each tube was incubated with functionalized NPs included in the Proteograph Assay Kit. A one-hour incubation with NP surfaces allowed for protein corona formation to reach equilibrium and was followed by a series of gentle washes using the super-paramagnetic properties of the NPs to remove non-specific and weakly bound proteins.

Proteins bound to the NPs were then reduced, alkylated, and digested with Trypsin/Lys-C to generate tryptic peptides for downstream LC-MS/MS analysis. All steps were performed in a one-pot reaction directly on the NPs. The in-solution digestion mixture was then desalted, and all detergents were removed using a mixed media filter plate and a positive pressure (MPE) system.

Clean peptides were then eluted in a high-organic buffer into a deep-well collection plate. Immediately after peptide elution, peptide quantitation assay was performed using the Pierce Fluorescent Assay Kit to determine the peptide yield for each well.

#### Sample Multiplexing with TMTpro 18-plex Labeling

As shown in Figure 2A, NP peptides were pooled, dried in a SpeedVac (3 hours), and then reconstituted directly in 50% acetonitrile in 100mM HEPES (pH 8) containing one of the TMT tags from the TMTpro 18-plex reagent (Thermo Fisher Scientific). The peptide-TMT mixture was incubated in a thermomixer for 1 h at 25°C and 600 rpm, and the reaction was stopped by addition of 2% hydroxylamine to a final concentration of 0.2% and incubated for 15 min at 25°C and 600 rpm. Labeled peptides for each 18-plex batch were pooled and dried using a SpeedVac system, and subsequently reconstituted in 0.1% FA for desalting using a C18 TopTip (PolyLC, Columbia, Maryland) according to the manufacturer’s recommendation.

**Figure 2:**
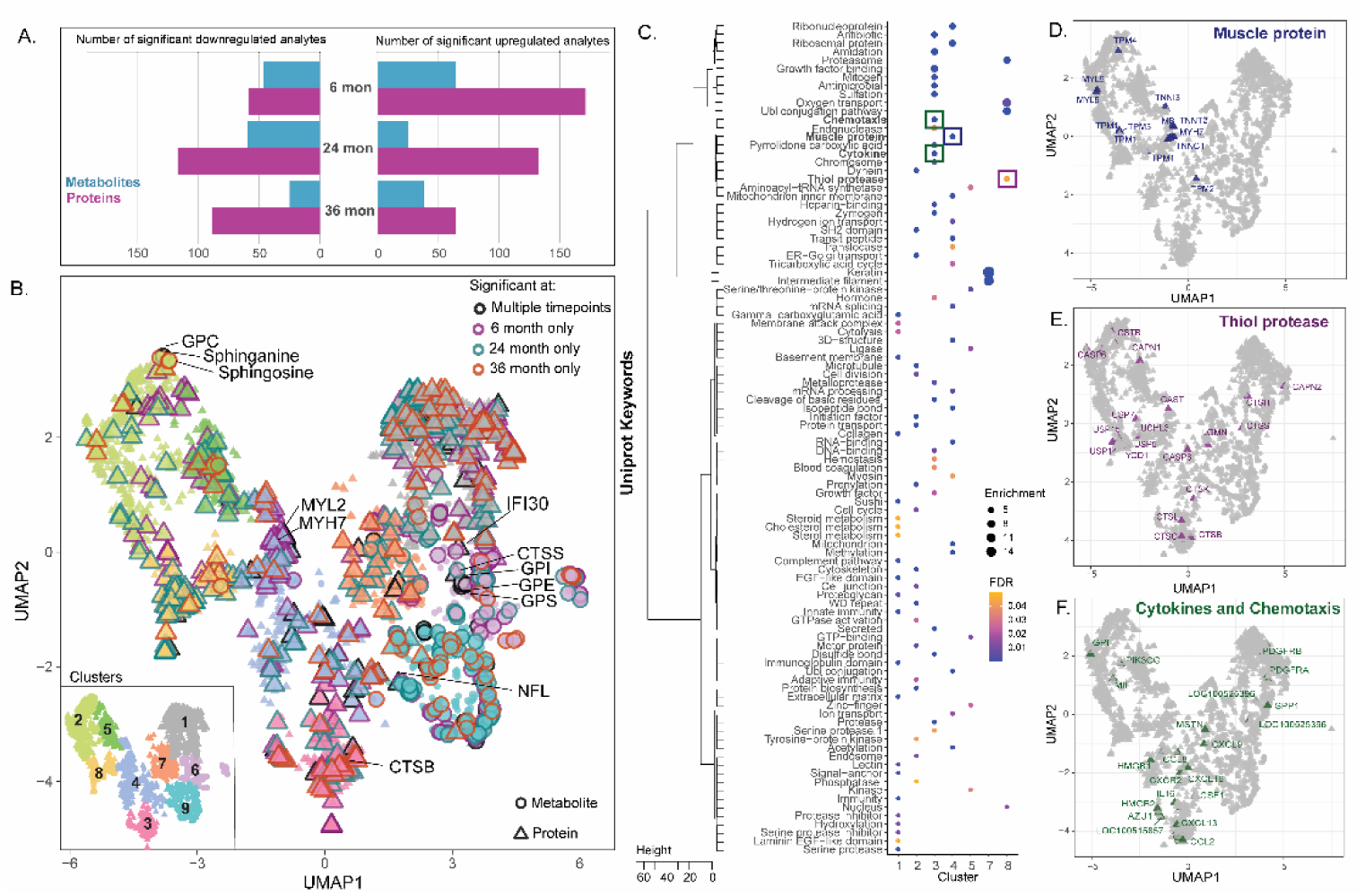
The Molecular Landscape of Longitudinal Metabolomics and Proteomics of the *CLN3* and Controls over Time. (A) Representation of the metabolites and proteins that significantly differentiate controls and *CLN3* at 6-, 24-, and 36-months. (B) UMAP of all metabolites (circles) and proteins (triangles) from all time points that has been divided into 9 clusters using hierarchical clustering from the Euclidean distance within the UMAP. Each cluster is represented by a different color (1 = grey, 2 = light green, 3 = pink, 4= light blue, 5 = green, 6 = light purple, 7 = orange, 8 = yellow and 9= light teal). The features that significantly differentiate controls from *CLN3* are outlined based on if they are significant at multiple time points (black), significant at the 6-month time point only (purple), 24-month time point only (teal), or significant at the 36-time point only (orange). Metabolites and proteins of interest are labeled with abbreviated names. (C) All proteins were mapped to their associated Uniprot Keywords. The clusters were tested for enrichment of keywords. The heatmap shows an overview of the enriched keywords in the clusters with enrichments greater than 3 and FDR corrected p-values less than 0.1, where the size of the circle is based on the enrichment and color is associated with the FDR corrected p-value. Enrichments of keywords are represented in (D) muscle protein (dark blue), (E) thiol protease (dark purple) and (F) cytokine/chemotaxis (dark green). Note that the “GPI” analyte in plot 2F refers to the protein glucose 6-phosphate isomerase (GPI) rather than the glycerophosphoinositol (GPI) metabolite discussed elsewhere.

#### Two Dimensional LC-MS/MS Proteomics Analysis

Desalted TMT-labeled peptide pools were dried in a SpeedVac system and reconstituted in 20 mM ammonium formate pH ∼10 for chromatography fractionation using a Waters Acquity BEH C18 column (2.1× 15 cm, 1.7 µm pore size) mounted on an M-Class Ultra Performance Liquid Chromatography (UPLC) system (Waters). Peptides were then separated using a 35-min gradient: 5% to 18% B in 3 min, 18% to 36% B in 20 min, 36% to 46% B in 2 min, 46% to 60% B in 5 min, and 60% to 70% B in 5 min (A=20 mM ammonium formate, pH 10; B = 100% ACN). A total of 72 fractions were collected and pooled in a non-contiguous manner into 36 total fractions. Pooled fractions were dried to completeness in a SpeedVac concentrator prior to mass spectrometry analysis.

Dried peptide fractions were reconstituted with 2% ACN, 0.1% FA and analyzed by Reversed Phase (RP) LC-MS/MS using an EASY-nLC^TM^ 1200 system (Thermo Fisher Scientific) coupled to an Orbitrap LumosTribrid^TM^ mass spectrometer equipped with FAIMS Pro^TM^ Interface (Thermo Fisher Scientific). Peptides were separated using an analytical C18 Aurora column (75µm x 250 mm, 1.6µm particles; IonOpticks) at a flow rate of 300 nL/min using a 80-min gradient: 1% to 6% B in 0.5 min, 6% to 23% B in 50 min, 23% to 34% B in 29 min, and 34% to 48% B in 0.50 min (A= FA 0.1%; B=80% ACN: 0.1% FA). The mass spectrometer was operated in positive data-dependent acquisition mode, and the FAIMS Pro Interface device was set to standard resolution with the temperature of FAIMS inner and outer electrodes set to 100°C. A three MS experiment method was set up where each experiment utilized different FAIMS compensation voltages: −45, −65, and −80 Volts, and each of the three experiments had a 1 second cycle time. A high resolution MS1 scan in the Orbitrap (m/z range 350 to 1,500, 60k resolution, AGC 4e5 with maximum injection time of 50 ms, RF lens 30%) was collected in top speed mode with 1-second cycle time for the survey and the MS/MS scans. For MS/MS (MS2) spectra, ions with charge state between +2 and +7 were isolated with the quadrupole mass filter using a 0.7 m/z isolation window, fragmented with higher-energy collisional dissociation (HCD) with normalized collision energy of 35% and the resulting fragments were detected in the Orbitrap at 50k resolution, at AGC of 5e4 and maximum injection time of 86ms. The dynamic exclusion was set to 20 sec with a 10 ppm mass tolerance around the precursor.

#### Proteomics Data Analysis

All mass spectra files were analyzed with SpectroMine software (Biognosys, version 2.7.210226.47784) using the TMTpro 18-plex default settings. The search criteria were set as follows: full tryptic specificity was required (cleavage after lysine or arginine residues unless followed by proline), 2 missed cleavages were allowed, carbamidomethylation (C), TMTpro (K and peptide n-terminus) were set as fixed modification and oxidation (M) as a variable modification. The false identification rate was set to 1% at peptide (or PSM) and protein levels. PSM report was exported from SpectroMine to R package tools for further analysis (data available in PX).

The R package {MSstatsTMT} version 2.2.7 was used to log_2_-transform the peptide intensities, impute within-TMT mixture missing values using an accelerated failure model, perform global median normalization on the peptide data (equalizing the medians across all channels and MS runs), conduct fraction aggregation, and perform protein quantification^18^. MSstatsTMT leverages a reference channel to perform local normalization, which effectively mitigates the systematic bias among different TMT mixtures. We conducted a comparison between utilizing the mean across all samples within each mixture as an artificial reference channel and using only the pooling of 48-month-old subjects as the reference channel. The results revealed that normalization based on all samples successfully minimizes between-mixture variance and eliminates the unwanted batch effect (Supplementary Fig. 2). Consequently, we adopted the normalization approach based on all samples^19^. Subsequently, we excluded the samples from 48-month-old subjects from downstream statistical analysis due to their limited biological variance. Finally, abundance values for each individual protein were scaled to have a mean of zero and unit variance. Proteins missing in 50% or more of the samples were excluded from the analysis, leaving 2,630 quantified proteins.

### Statistical and Enrichment Analysis

Analytes (proteins and metabolites) that differed in abundance between the wild type and *CLN3*^Δex7–8^ samples at each time point (6-, 24-, or 36-months) were identified via two-tailed Student’s t-test. Analytes with an uncorrected p-value < 0.05 were associated with genotype. The sets of proteins associated with genotype at each time point were assessed for enrichment in GO terms using Database for Annotation, Visualization, and Integrated Discovery (DAVID). The sets of metabolites associated with genotype at each time point were assessed for enrichment in chemical structure types using MetaboAnalyst^20^. In both cases, Fisher’s exact test was performed with Benjamini-Hochberg false discovery rate correction, and all detected proteins or metabolites were used as the background in the enrichment analyses, as applicable.

Datasets for the proteins and metabolites that were normalized and scaled were combined for a final dataset. The final dataset was reduced to a 2-dimensional uniform manifold approximation and projection (UMAP) using the uwot package in R^21^. The UMAP was then clustered using hclust (stats package in R) and the distance provided by the UMAP into 9 clusters, where the number of clusters was decided by the elbow method. The proteins from each cluster were then tested for enrichment of Uniprot Keywords using the AnnoCrawler pipeline (https://github.com/DansenCode/AnnoCrawler). Significant enrichment was determined by an enrichment of a keyword in comparison to the background (the rest of the UMAP) with an FDR cutoff of 0.1.

#### Multiblock Sparse Partial Least Squares Discriminant Analysis

Multiblock sparse partial least squares discriminant analysis (sPLS-DA), also known as DIABLO (Data Integration Analysis for Biomarker discovery using Latent component method, was performed on the processed proteomics and metabolomics datasets using the {mixOmics} package in R^22, 23^. This technique finds a linear combination of input variables from two or more omics datasets that reduces the dimensionality of the datasets while maximizing the covariance of the reduced datasets with each other and with an outcome variable. Specifically, in the case of a centered and scaled metabolomics dataset *X*(*M*)(N x P_1_) and centered and scaled proteomics dataset *X*(*P*)(N x P2) with genotype labels *Y* (N x G), where N is the number of samples, P1 is the number of metabolites, P^2^ is the number of proteins, and G is the number of groups, for each dimension ℎ = 1, …, *H* multiblock sPLS-DA finds the loading vectors 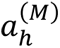 and 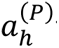 that maximize the value of:

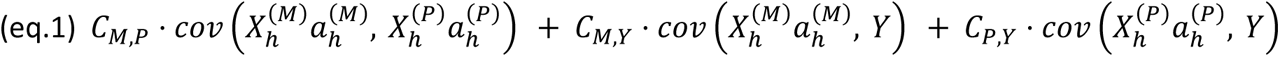

subject to 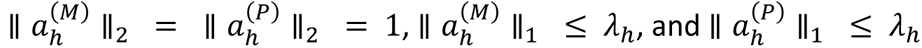

The values of the constants *C*_*M*,*P*_ *C*_*M*,*Y*_, and *C*_*P*,*Y*_ are chosen to reflect the expected degree of association between the different -omics datasets and between the -omics datasets and genotype. In this case, all constants were set to one (*i.e.,* a fully connected design matrix was used). The value of *λ*_ℎ_ is chosen to constrain the number of non-zero elements in the loading vectors 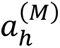 and 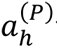

In this way, multiblock sPLS-DA selects a subset of variables (those with non-zero coefficients in 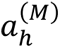and 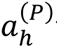) that are correlated both within and between the proteomics and metabolomics datasets and that define, for each omics dataset *X*^(*O*)^ and each dimension ℎ, the component score:

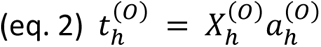

The process is iterative, such that the loading vector for the first dimension, 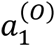 is found by maximizing equation 1 with 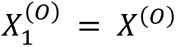, while the loading vector for the second dimension, 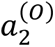, is found by maximizing equation 1 with 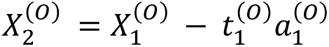, and so on for however many dimensions are desired. This ensures that the first components 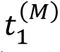 and 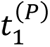encompass the greatest variation in the dataset, followed by the second components 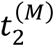 and 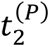, etc.

The implementation of multiblock sPLS-DA in the {mixOmics} package creates a classifier that calculates the components 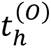 for a new sample, makes one classification for that sample per omics dataset in the model (based on the distance between the component scores for the new sample and the stored component scores for the training dataset points), then makes a final classification based on either majority vote, weighted vote, or averaged vote of the individual omics classifiers.

Because we desired a continuous disease score that incorporated both the proteomics and metabolomics datasets in our study, rather than the binary classifier created by default in mixOmics, we created an “sPLS score” by summing together the first component scores for the metabolite and protein datasets.

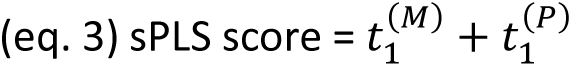

We used this sPLS score to assess the combined ability of the proteins and metabolites selected in the loading vectors to distinguish *CLN3^Δex7–8^* from wild type samples in a held-out test set.

Prior to performing multiblock sPLS-DA, the data was randomly split into a training set (70%, n=34) and test set (30%, n=14). No individual subject was represented in both the training and test sets (*i.e.*, if a subject was sampled at multiple time points, the data for both time points were included together in either the training or test set). Multiblock sPLS-DA was performed on the training set using two dimensions (h=1,2). The number of non-zero coefficients to use in the loading vectors was determined by assessing the balanced error rate of the mixOmics-generated classifier on the training dataset with five-fold cross-validation repeated 20 times. The lowest average balanced error rate (0.004 ± 0.010) was achieved by using one protein in the protein first component and three metabolites in the metabolite first component. However, we chose to include two proteins in the protein first component (two non-zero coefficients in 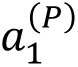) and two metabolites in the metabolite first component (two non-zero coefficients in 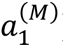) because this also yielded a very low average balanced error rate (0.026 ± 0.023) and presented the benefit of drawing two analytes from each of our omics datasets. A single analyte was used for each second component (one non-zero coefficient each in 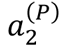 and 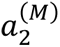) because the number of analytes selected for the second components had little impact on classification accuracy, and because the second components tended to correlate with age rather than genotype.

For the proteins, the first component was initially a linear combination of the normalized abundance values for CTSS and ADAMTSL4. However, due to lack of availability of commercial kits for testing ADAMTSL4 levels in patient samples, we removed ADAMTSL4 from consideration and repeated the multiblock sPLS-DA workflow.

## Results

To investigate longitudinal blood-based CLN3 disease signatures, we performed deep multi-omics profiling to quantify relative blood serum concentrations of 769 metabolites and 2,634 proteins in samples from male and female wild type and homozygous *CLN3^Δex^*^7–8^ Yucatan Minipigs. 6-, 24-, and 36-month time points were selected representing early-stage disease prior to any appreciable visual or motor defects, mid-stage/symptomatic, and late-stage disease, respectively (Fig. 1 and Supplementary Fig. 3). Of the 3,403 analytes quantified in the dataset, 230 proteins (171 upregulated; 59 downregulated) and 111 (64 upregulated; 47 downregulated) metabolites were associated with genotype at the 6-month time point, suggesting early systemic changes. At 24-months, 249 proteins (132 upregulated; 117 downregulated) and 85 metabolites (25 upregulated; 60 downregulated) were associated with genotype, while at the 36-month timepoint 153 proteins (64 upregulated; 89 downregulated) and 63 metabolites (38 upregulated; 25 downregulated) were associated with genotype (Fig. 2A). To discern large-scale relationships among the combined metabolomics and proteomics datasets, we employed the Uniform Manifold Approximation and Projection (UMAP) algorithm to reduce the data into a 2-dimentional space (Fig. 2B). The UMAP algorithm was used because it is effective in visualizing clustering patterns in high-dimensional data, therefore the distance between the features is relative to the way the analytes co-vary in the dataset and associate in the 2-dimentional space. The features are colored by cluster and have bold outlines if they were found to be significant at one or more time points. The features in the UMAP were separated into 9 distinct clusters, each enriched for certain Uniprot Keywords (Fig. 2C). Shown are enrichments greater than 3 with an FDR corrected p-value less than 0.05, with the full list in Supplementary Table 1. Specific Uniprot keywords (Muscle protein, Thiol protease, Cytokine/Chemotaxis) were selected to show the enrichment throughout the UMAP of selected relevant biological features (Fig. 2D-F). Consistent with previous studies, we found a group of structurally similar glycerophosphodiesters (GPI, GPE, GPS; Fig. 2B) among the most significantly and consistently upregulated species across all time points, while a group of phospholipids sharing a docosahexaenoyl group at the sn2 position were among the most significantly and consistently downregulated analytes^8^.

In addition to these previously identified analytes, deep nanoparticle-based proteomics workflow (Fig. 3A) combined with isobaric labeling and 18xmultiplexing (TMT) quantified over 2600 proteins, several of which were strongly associated with Batten disease genotype (Fig. 3B-G). The top two differentially expressed proteins across all time points were Cathepsin S (CTSS) and Cathepsin B (CTSB), lysosomal cysteine proteases that both demonstrated remarkably stable longitudinal patterns of elevation (Fig. 3B, C). Although not significant at all time points, gamma-interferon inducible lysosomal thiol reductase IFI30 (IFI30), Myosin regulatory light chain 2 (MYL2), and Myosin-7 were also elevated in *CLN3* mutant samples (Fig. 3D-F). As previous studies have shown, neurofilament light (NFL) was able to significantly differentiate controls and CLN3 disease but only at 36 months (Fig. 3G). Using the novel streamlined nanoparticle-based proteomics workflow provided several advantages. In contrast to targeted affinity-based approaches that require species specific affinity probes, the nanoparticle workflow is directly amenable to non-human model organisms. Moreover, direct compatibility with multiplexing at the peptide level enhanced throughput and data robustness. Together, this enabled the deepest Batten disease as well as porcine serum proteome profile to date, quantifying novel putative disease biomarkers commonly not detectable with standard serum proteomics workflows (Fig. 3H and Supplementary Table 2).

**Figure 3:**
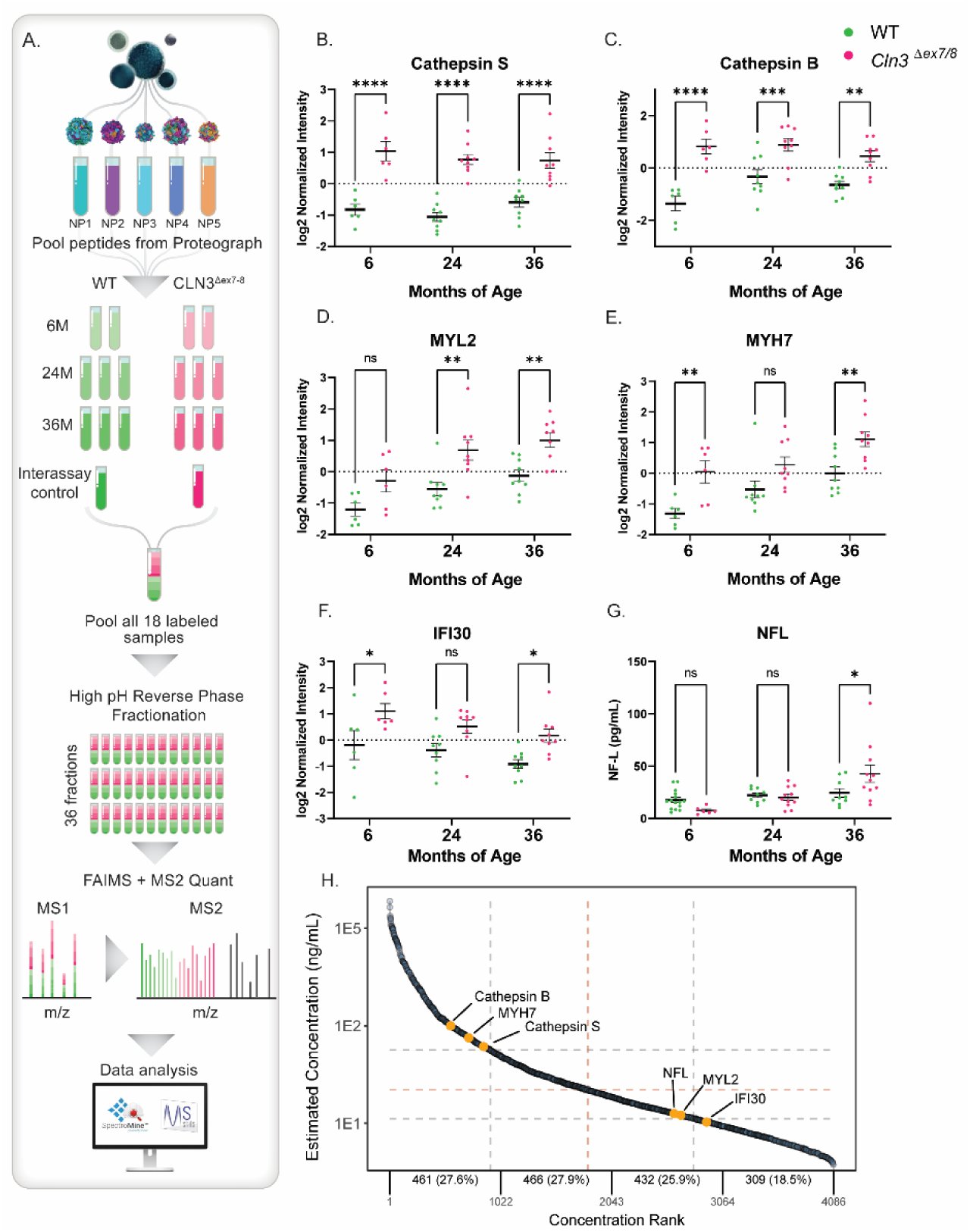
Proteograph Workflow with TMT 18-plex Mass Spectrometry Method and Resulting Protein Biomarker Regulations. (A) Serum samples were first processed by SP100 Automation instrument with ProteographTM Assay kit included in Proteograph Product Suite (Seer, Inc.) using five distinctly functionalized nanoparticles (NPs). Tryptic peptides from 5 NPs were then pooled into one single sample for TMT labeling. A total of 54 pooled samples were allocated into three 18-plex TMT mixtures, each of which contained two 6-month samples, three 24-month samples, three 36-month samples and one interassay control sample from WT and *CLN3^Δex^*^7–8^ respectively. Each TMT mixture was followed by high pH reverse phase fractionation and LC-MS/MS analysis, comprised of a Proxeon EASY nanoLC system coupled to an Orbitrap Fusion Lumos MS equipped with FAIMS Pro Interface (Thermo Fisher Scientific). The raw spectra data were finally processed by SpectroMine (Biognosys) and R/Bioconductor package MSstatsTMT to generate statistical analysis results. (B-F) Lysosomal proteins Cathepsin S (CTSS), Cathepsin (CTSB), and IFI30 and muscular proteins MYL2 and MHY7 are significantly elevated at multiple time points in *CLN3^Δex^*^7–8^ minipig serum when compared to age-matched healthy control animals, (G) while NFL blood-serum levels are significantly elevated at 36 months only. Two-way ANOVA with Šidák correction for multiple comparisons, 95% confidence interval, *p<0.05, **p<0.01, ***p<0.001, ****p<0.0001, n = 6; 9; 9 animals respectively, mean +/-SEM. (H) The 2,634 identified proteins were mapped to the HPPP protein database, revealing their detection throughout the entire concentration range of the database. Notably, the proteins NFL, MYL2, and IFI30 were found to be within the low abundance range.

To elucidate the functional relevance of the differentially expressed proteins, we probed for enriched functional sets of proteins and pathways within the subsets of proteins associated with genotype at each time point. Three ‘cellular component’ signatures were flagged at all time points: cytoplasm, proteasome, and secreted (Supplementary Table 3). Interestingly, the signature “lysosome” was shared only between the 24-and 36-month groups suggesting that blood signatures of lysosomal dysfunction increase over time, perhaps reflecting progressive dysfunction in this organelle. When categorized by “biological process,” the related term “chemotaxis” was shared among all groups, suggesting alterations in immune homeostasis. (Supplementary Table 4). Unique to the 36-month time point, we found a large network of proteins and metabolites involved in sphingolipid metabolism. Characterized by a common eighteen carbon amino-alcohol backbone, sphingolipids are crucial to a vast number of biological processes including cell signaling and membrane structure and have previously been linked to CLN3 Batten disease, establishing another putative functional link between biomarker signatures and pathogenicity^24–26^.

Kyoto Encyclopedia of Genes and Genomes (KEGG) pathway analysis confirmed the enrichment for species involved in sphingolipid metabolism at 36 months. We detected the presence of 21 analytes with known function in sphingolipid metabolism and related pathways, 11 of which were differentially abundant in *CLN3* serum (Fig. 4 and Supplementary Fig. 4). Notably, all the differentially abundant metabolites were elevated in *CLN3* serum, suggesting alterations in sphingolipid synthesis, degradation, or defective transport of intermediates (Fig. 4 and Supplementary Fig. 4A-F). Many proteins contributing to sphingolipid metabolism were similarly elevated, suggestive of a compensatory mechanism triggered by the accumulation of sphingolipids and resulting increased lysosomal burden (Fig. 3H-M and Supplementary Fig. 4G-L). Given the complex nature of this pathway, further investigation is needed to clarify the impact of CLN3 Batten disease on sphingolipid metabolism, or vice versa.

**Figure 4:**
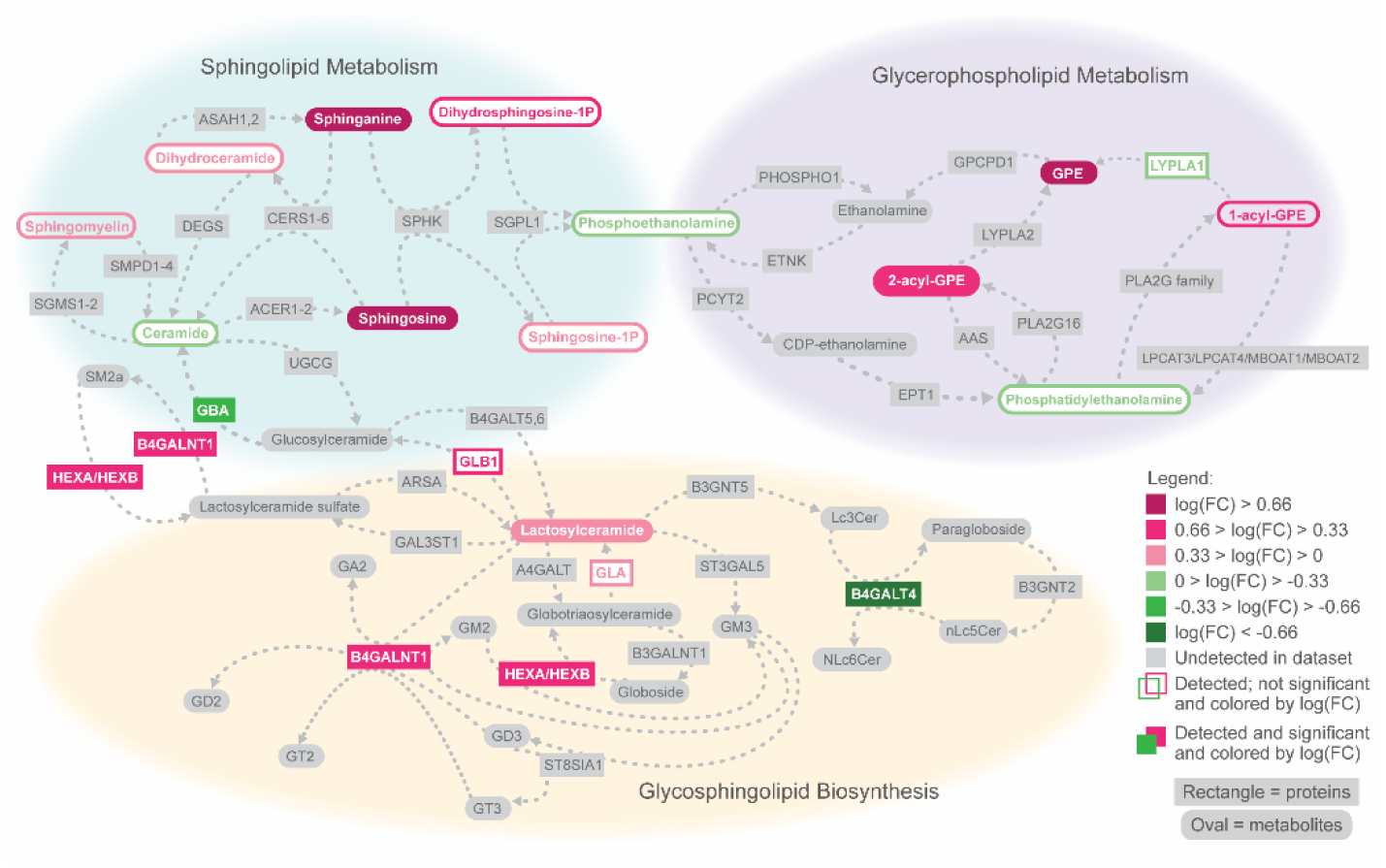
Sphingolipid Alterations at 36 Months in CLN3 Disease. Network of proteins and metabolites involved in sphingolipid metabolism and related pathways that were differentially abundant in serum of 36-month-old subjects. Network connections were derived from the KEGG pathways for sphingolipid metabolism (top left), glycerophospholipid metabolism (top right), and glycosphingolipid biosynthesis (bottom). Serum levels of B4GALT4 and GBA were significantly lower in *CLN3*^Δex7–8^ pigs than in WT pigs, while serum levels were significantly higher for B4GALNT1, HEXA, HEXB, lactosylceramide, sphinganine, sphingosine, glycerophosphoethanolamine, sphingomyelin, and 1-acyl-GPE (uncorrected Student’s t-test, p < 0.05). Analytes not significantly different between genotypes are outlined according to the magnitude and direction of their log fold change, while analytes involved in the pathway that were not detected in the dataset are presented in gray. Many of the metabolites in the KEGG pathways are general metabolite categories rather than specific chemical species (e.g., ceramide, 1-acyl-GPE). Multiple analytes were detected in the dataset that fall under the categories of 1-acyl-GPE (n=5), ceramide (n=6), phosphatidylethanolamine (n=10), and sphingomyelin (n=28). Not all detected analytes within these categories behaved the same way (supplementary information); the color of the category in the figure is based on the overall trend. Other categories had only one representative detected in the dataset, including 2-acyl-GPE (2-stearoyl-GPE (18:0)), dihydroceramide (N-palmitoyl-sphinganine (d18:0/16:0)), and lactosylceramide (lactosyl-N-palmitoyl-sphingosine (d18:1/16:0)).

We investigated biomarkers manifesting early in disease progression, prior to the onset of symptoms which could have utility for pre-symptomatic clinical diagnosis, early evaluation of interventions, and providing insights into upstream disease processes (Supplementary Fig. 5). Of the 230 genotype-associated protein targets at 6 months, Integrin Beta-2 (ITGB2) was among the most significantly downregulated protein, although ITGB2 serum levels rebounded at later time points, remaining unchanged at 24 and 36 months (Supplementary Fig. 5A). Conversely, Calpastatin (CAST) and Myosin Light Chain 3 (MYL3) – cardiac-enriched proteins - were significantly elevated in the *CLN3* cohort at 6 months when compared to wild type, with smaller elevations at later time points (Supplementary Fig. 5B, C). Glucuronide of C12H2OO3 and 3-aminoisobutyrate were downregulated at 6 months with a rebound at later time points (Supplementary Fig. 5D, E). Interestingly, maleate – showing the largest fold change of any upregulated metabolite species at 6 months – stabilizes to near identical levels in *CLN3^Δex^*^7–8^ and wild type minipig serum at 24- and 36-months (Supplementary Fig. 5F). Together these data demonstrate complex pre-symptomatic molecular changes at the protein, lipid and metabolite levels throughout disease progression that could serve to assess therapeutic efficacy.

Given the immense scale and complexity of the multi-omics dataset, we sought to condense these data into a disease score model that could reflect overall disease signatures derived from a minimal set of input variables. Similar to tools such as multi-domain responder indices that have found favor in clinical trials, multiblock sPLS-DA was used to reduce the proteomics and metabolomics training datasets into two components each. For the protein detectable with the nanoparticle-based workflow, the first component comprised CTSS and CTSB (Fig. 5A). For the metabolites, the first component was a linear combination of the normalized abundance values for glycerophosphoinositol (GPI) and glycerophosphoethanolamine (GPE) (Fig. 5B). For both datasets, the first component separated the samples by genotype, while the second component separated the samples by age (Fig. 5A, B).

**Figure 5:**
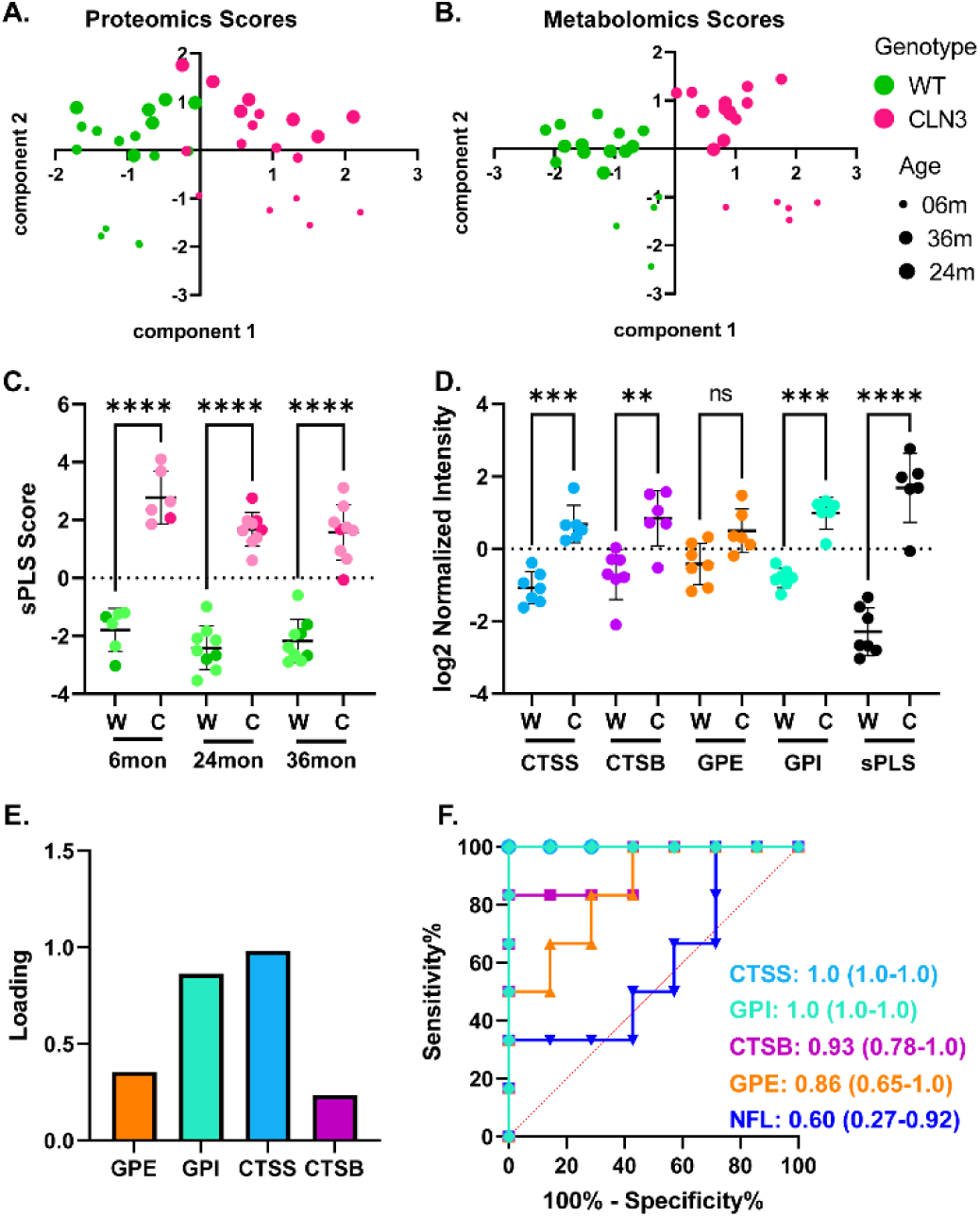
Multiblock sPLS-DA Loadings Highlighting Protein Biomarkers for *CLN3*. A,B. Training data projected onto the two proteomics components and two metabolomics components identified using multiblock sPLS-DA. For both omics datasets, the first component separated the datapoints by genotype, and the second component separated the datapoints by age. C. sPLS scores calculated for the entire dataset, with test set points shown in dark green/magenta. At each time point, sPLS scores were significantly different between WT and *CLN3^Δex^*^7–8^ subjects in the dataset as a whole. Scores were also significantly different between WT and *CLN3^Δex^*^7–8^ subjects in the test set at all time points combined (p < 0.0001, Student’s two-tailed t-test), demonstrating the robustness of the sPLS score as a tool for differentiating *CLN3^Δex^*^7–8^ from WT samples. D. Comparison of sPLS score and log2 normalized intensity values of the analytes that make up the sPLS score for separating data in the test set. The sPLS score better separates the test set WT from *CLN3^Δex^*^7–8^ samples compared to any individual analyte (bars represent standard deviation; asterisks represent p-values from two-way ANOVA with Tukey’s post-hoc test). E. Loadings of the analytes in the sPLS score created by combining the formulas for the protein first component and metabolite first component. The formula for the protein first component relied on cathepsin S (CTSS) and cathepsin B (CTSB), while the formula for the metabolite first component relied on glycerophosphoinositol (GPI) and glycerophosphoethanolamine (GPE). The sPLS score formula places the greatest emphasis on CTSS, followed by GPI, then GPE, and finally CTSB. F. Receiver operating characteristic curves for each of the analytes included in the sPLS score, as well as neurofilament light (NFL), as classifiers for identifying *CLN3^Δex^*^7–8^ samples in the test set. The area under the ROC curve is presented for each analyte along with its 95% confidence interval (computed with Wilson/Brown method). CTSS and GPI both serve as perfect classifiers for the test set, and all four analytes included in the sPLS score outperform NFL as individual classifiers.

The predictive accuracy of the model was assessed with a held-out set of test samples, where sPLS scores were significantly different between *CLN3^Δ^*^ex7–8^ and wild type samples (two-tailed t-test, p < 0.0001) from all three time points (Fig. 5 C, D). The sPLS score served as a perfect classifier for the test set (ROC AUC=1.0, Supplementary Information) and separated the wild type and *CLN3^Δ^*^ex7–8^ samples in the test set better than any individual analyte used to construct the sPLS score (Fig. 5D). The weight for each of the selected analytes in the model can be seen in the loadings plot (Fig. 5E). The ROC curves (Fig. 5F) validate these findings by showing perfect separation for CTSS and GPI (ROC AUC =1.0), whereas CTSB (AUC = 0.93) and GPE (AUC = 0.86) are better at determining Batten Disease as compared to NFL (AUC = 0.6).

## Discussion

The absence of functional CLN3 gives rise to an intricate disease state commonly referred to as Batten disease. Cells lacking functional CLN3 protein exhibit a wide range of abnormalities including trafficking deficits within the secretory pathway, changes in lipid composition, impaired autophagy, and altered lysosomal composition and function^26–29^. At the tissue level, the central nervous system is progressively impaired by extensive neuroinflammation^30^, while there is some evidence of progressive dysfunction in the heart, skeletal muscles, and immune cell populations in the periphery^31^. Despite comprehensive characterization of disease signs and symptoms, there is little consensus around the primary, secondary, or tertiary etiologies emerging from CLN3 dysfunction. Furthermore, there is a notable dearth of information on clinically measurable biomarkers that could be used to monitor disease progression or therapeutic efficacy.

Our study addresses these knowledge gaps with longitudinal deep multi-omics profiling using blood serum from a large animal model of CLN3 disease, the *CLN3^Δex7/8^* minipig. The *CLN3^Δex7/8^* minipig exhibits progressive disease pathology and behavioral abnormalities, more closely resembling human disease as compared to currently available small animal models such as the *Cln3^Δex7/8^* mouse. Additionally, the use of *CLN3^Δex7/8^* minipig in biomarker studies offers a controlled, isogenic, and well-powered approach compared to the challenges of studying rare samples from individuals with CLN3 disease with wide genetic variation. A particular challenge with animal models is the extraordinarily large dynamic range of protein concentrations in blood and lack of appropriate affinity probes to either deplete or enrich proteins to facilitate comprehensive proteome capture at scale. Our study demonstrates the utility of a novel nanoparticle-based proteomics workflow that mitigates the dynamic range challenges in a species-agnostic way to facilitate unbiased and deep serum proteomics. In combination with global metabolomic and lipidomic profiling, we quantified over 3400 analytes longitudinally (2,634 proteins; 769 metabolites and lipids), significantly surpassing previous porcine multi-omic studies, which were hindered by the limitations of human- or mouse-specific depletion kits^32, 33^. The combination of the Proteograph workflow with TMTpro 18-plex reagents has enabled the simultaneous analysis of 18 samples, increasing the throughput and accuracy of protein quantification with enhanced data completeness^34^. Further, our workflow for multiplexed proteomics data acquisition using MS2 methods on an Orbitrap Tribrid MS instrument utilizing the FAIMS Pro interface has shown to enhance quantification accuracy^35^.

With this unique approach, we were able to identify biomarker signatures that offer new insights into the cellular dysfunction and tissue pathology associated with the loss of functional CLN3, as well as a more detailed understanding of the disease timeline. The early emergence and steady maintenance of elevated glycerophosphodiesters suggests a close association with CLN3 function, while later in the disease course a second signature comprised of sphingolipid metabolism proteins and metabolites emerges. A persistent lysosomal signature is also present throughout disease progression, likely due to increased lysosomal mass (from storage material accumulation and upregulation of lysosomal biogenesis) either exocytosed or released into circulation by progressive inflammation and tissue damage.

Although the role of glycerphophodiesters is not completely understood in Batten disease, elevated levels have been observed early in the course of the disease and are maintained at steady levels, suggesting a close association with CLN3 function. One likely possibility is that glycerophosphodiesters are an early lysosomal storage substrate, leading to subsequent secondary storage of additional proteins, lipids, and metabolites. If this is the case, targeting the upstream metabolic pathways related to glycerophosphodiester metabolism may have therapeutic potential. Later in disease progression, effective treatments may need to target additional facets of disease such as immune activation, aberrant sphingolipid metabolism, and downstream lysosomal pathology. Importantly, our data suggests that some of the molecular patterns are temporally restricted, suggesting that an ideal therapeutic strategy could potentially be tailored to an individual’s specific biomarker signature of disease progression.

We discovered several proteins, many of which would be hard or impossible to detect with conventional unbiased proteomics workflows at scale (Supplementary Table 2), exhibiting differential expression between CLN3 Batten disease and control samples. Among them, lysosome associated CTSS and CTSB were consistently upregulated at all time points which is consistent with the central role of lysosomal dysfunction in the disease pathology. Another lysosomal protein, IFI30, exhibited similar elevations, although differences did not reach significance at the six-month time point. Why these lysosomal proteases are enriched in a peripheral biofluid remains unclear. One likely possibility is that CTSS, CTSB, and IFI30 are enriched in disease-affected cells, possibly as a consequence of increased lysosomal mass or as a compensatory mechanism for clearing storage material. The proteins could then be released into circulation upon lysosomal exocytosis or cell death. The cardiac-enriched proteins MYL2 and MYH7 showed similar patterns of elevation, reaching significance at most time points. It is tempting to speculate that these markers could reflect a cardiac defect in this model – a hypothesis that warrants further investigation. Cardiac phenotypes (left ventricular hypertrophy, bradycardia, storage material) have been described in CLN3 affected individuals, but a lack of strong phenotypes in mouse models has thus far precluded exhaustive preclinical study of this facet of disease^36^.

These new protein-based biomarkers could be valuable additions to the repertoire of existing markers for CLN3 disease. One existing marker, NFL, offers the benefit of being closely linked to the central neuronal pathology of the disease, but exhibits only mild elevations in affected individuals and animal models, as reflected in our data here. The glycerophosphodiesters are appealing biomarker candidates due to their early and steady elevation, but little is known about their relationship with disease etiology. In combination, however, CTSS, CTSB, GPI, and GPE offer a powerful summary of disease state from a limited set of variables, as evidenced by our sPLS score model. Further development of a CLN3 disease biomarker panel could also incorporate markers of neuronal damage, such as NFL, to build an even more comprehensive picture of an individual’s disease status.

Our data also provides new insights into the pathogenic timeline of Batten disease. While CTSS, CTSB, GPI, and GPE were elevated at all time points, other signatures developed only at later stages of disease progression. For example, MYL2 and NFL were significantly upregulated only in the later time points, suggesting that cardiac and neuronal damage may take substantial time to initiate. Additionally, sphingolipid metabolism was only enriched at the 36-month time point, suggesting that sphingolipid accumulation takes longer to develop and may contribute to disease pathology in a more tissue specific manner before it is a useful biomarker in blood. Previous studies have similarly found sphingolipid metabolism to be dysregulated in various models of CLN3 disease, although exactly how this process contributes to disease remains unclear^37, 38^. Regardless of the mechanism, treating these secondary or tertiary disease manifestations with targeted therapies may be beneficial alongside disease-modifying therapies that compensate for the loss of CLN3.

Collectively, this work demonstrates how deep multi-omics profiling can be employed to detect disease-specific biomarkers related to cellular dysfunction and tissue pathology associated with a complex degenerative disease. Innovative techniques to mitigate interference of highly abundant proteins in pig serum were crucial in enabling comprehensive proteomics. The power of this technique is evidenced not only by the identification of proteins and metabolites as biomarker candidates, but also by the insights into the timeline and order of disease progression. This study’s findings demonstrate the potential of deep multi-omics profiling for uncovering disease-specific biomarkers, which can provide valuable insights for understanding disease mechanisms and identifying potential drug targets.

---

**Supplementary Figure 1:**
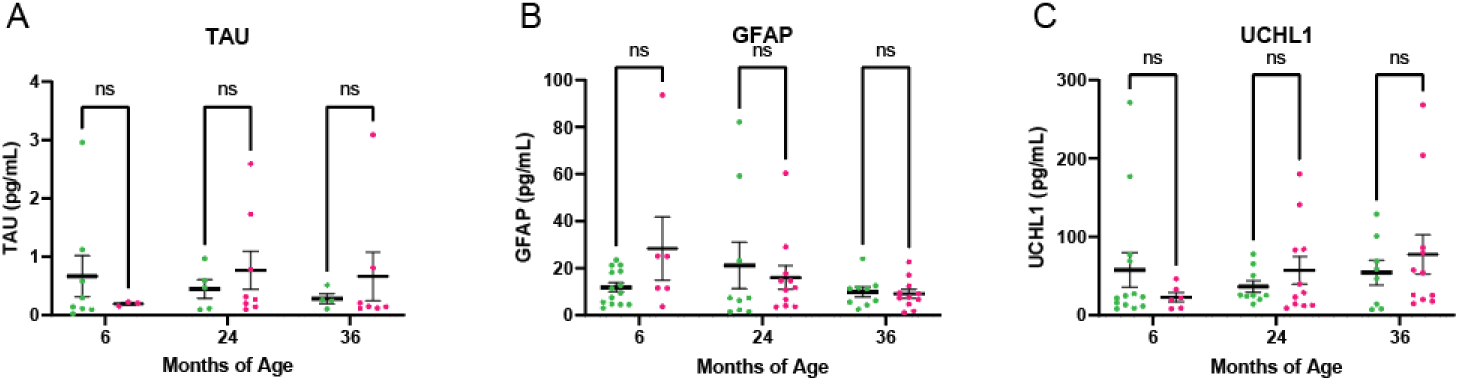
Neurology 4-PlexA Targeted Proteomic Analysis. The total measurement of (A) TAU, (B) glial fibrillary acidic protein (GFAP), (C) and ubiquitin carboxyl-terminal hydrolase L1 (UCHL1) in wildtype *CLN3^Δex7/8^* minipig serum at 6-, 24- and 36-months. No significant differences were observed. Two-way ANOVA with Šidák correction for multiple comparisons, 95% confidence interval, n = 6; 9; 9 animals respectively, mean +/-SEM.

**Supplementary Figure 2:**
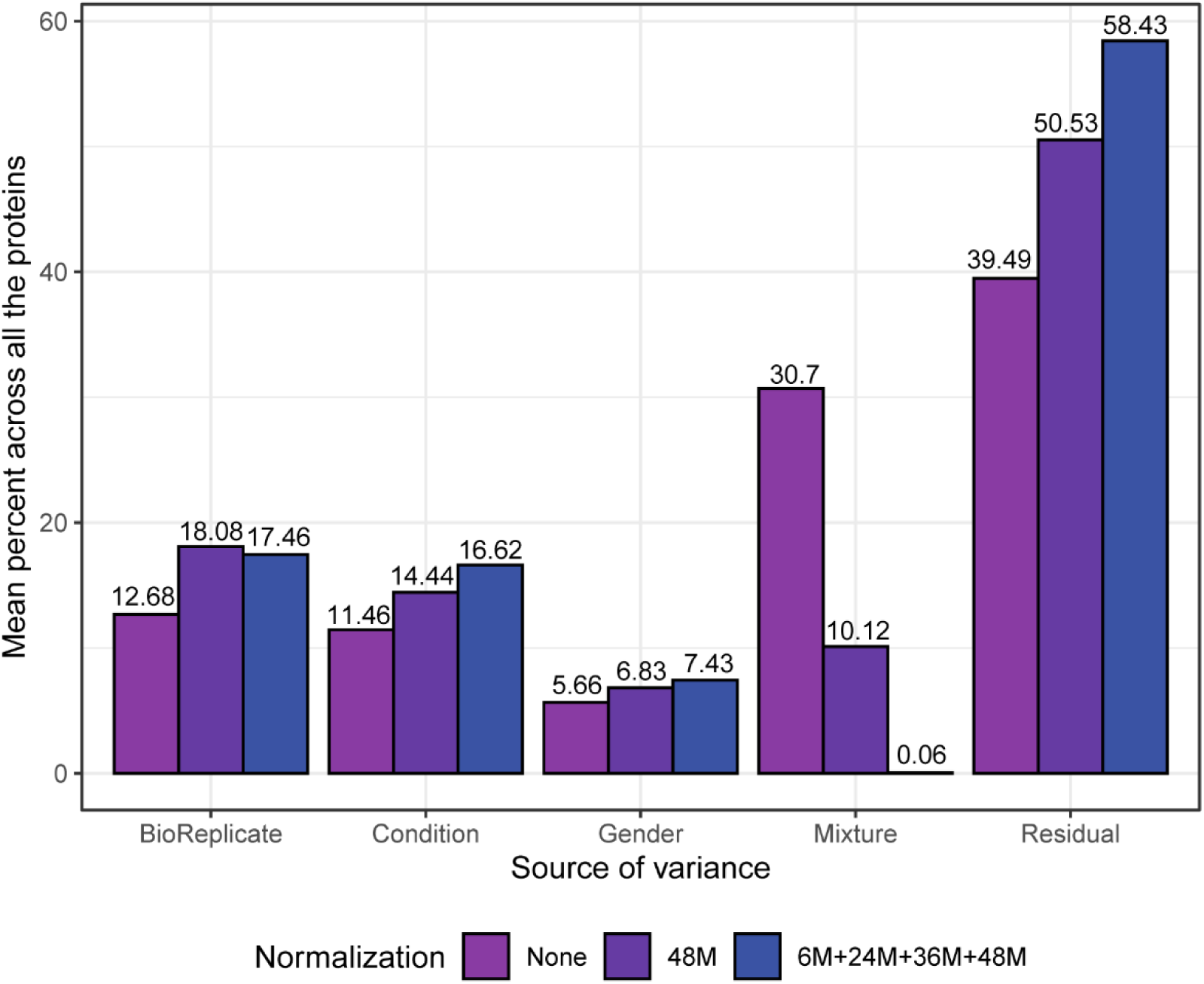
Variance Components Analysis on the Protein Intensities Generated by Different Normalization Methods. In the context of normalization methods, “None” indicates no normalization was performed. “48M” refers to the usage of pooling samples from 48-month-old subjects as the reference channel, while “6M+24M+36M+48M” signifies the utilization of the mean across all samples from all ages as the artificial reference channel. The x-axis represents the sources of different variance components, with “Conditions” representing the combination of age and genotype, and “BioReplicate” representing distinct pigs. The percentage of each variance component in the overall variation was calculated for each protein. The y-axis displays the average percentage of each variance component across all proteins. Utilizing normalization based on all the samples effectively mitigates the between-mixture variance, eliminating the undesirable batch effect.

**Supplementary Figure 3:**
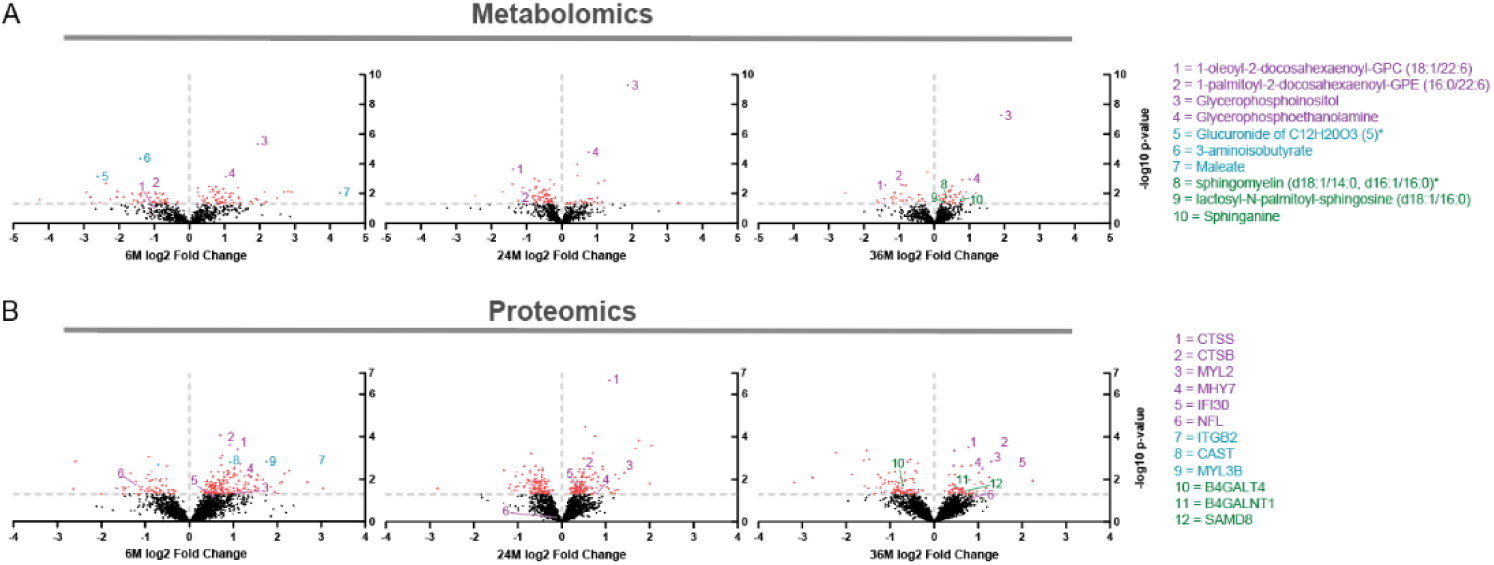
Profiling Differentiated Metabolites. (A) Volcano plots of detected metabolites with differentially expression levels identified at 6-, 24-, and 36-months (-log10 p-value > 1.3) highlighted in red. Top metabolite biomarker candidates across all time points denoted with purple, early biomarker candidates denoted by teal, and metabolites involved in sphingolipid metabolism are highlighted green. (B) Volcano plots of detected proteins with differentially expressed proteins at 6-, 24-, and 36-months (-log10 p-value > 1.3) highlighted red. Top protein biomarker candidates across all time points denoted by purple, early biomarker candidates denoted by teal, and proteins involved in sphingolipid metabolism are highlighted green. Student’s t-test, n = 6; 9; 9 animals respectively.

**Supplementary Figure 4:**
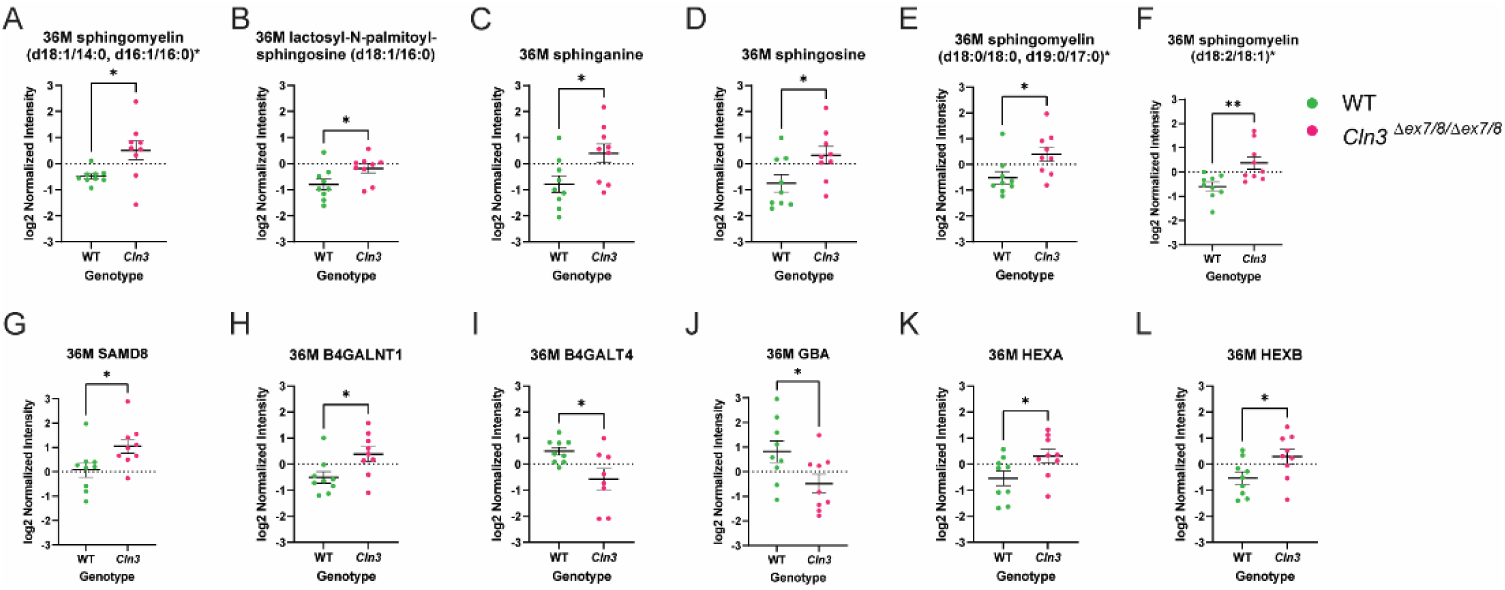
Metabolites and Proteins Involved in Sphingolipid Metabolism. Metabolites and proteins involved in sphingolipid metabolism were compared in wildtype and *CLN3^Δex7/8^* minipig serum samples at 36-months. (A-F) Differentially expressed metabolites at 36-months included sphingomyelin (d18:1/14:0, d16:1/16:0)*, lactosyl-N-palmitoyl-sphingosine (d18:1/16:0), sphinganine, sphingosine, sphingomyelin (d18:0/18:0, d19:0/17:0)*, and sphingomyelin (d18:2/18:1)*, all of which were upregulated in *CLN3^Δex7/8^* serum samples. (G-L) Sphingolipid proteins SAMD8, B4GALNT1, HEXA, and HEXB were significantly upregulated in *CLN3^Δex7/8^* minipigs, while B4GALT4 and GBA were significantly downregulated.

**Supplementary Figure 5:**
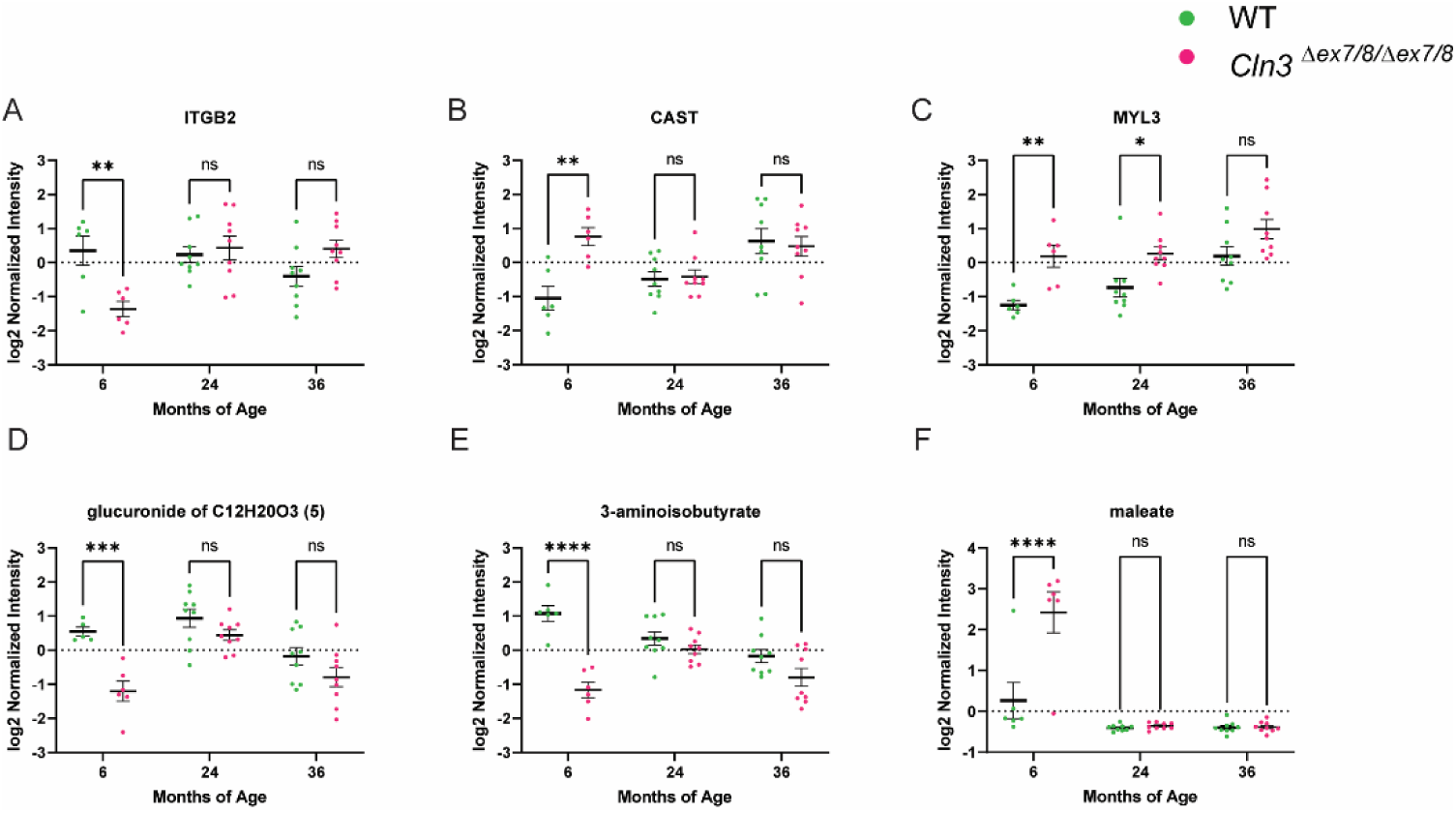
Identification of Biomarkers Appearing Early in Disease Progression, Prior to the Onset of Symptoms. (A-C)) Of the 230 protein targets associated with genotype at 6-months, ITGB2, CAST, and MYL3 demonstrate significant dysregulation, all stabilizing to wild type levels at later time points. (D-F) Similarly, Glucuronide of C12H2OO3, 3-aminoisobutyrate, and maleate demonstrate utility as “early” metabolite biomarkers and are only dysregulated at 6-months.

**Supplementary Table 1:** Can be found in the Excel sheet <u>here</u>.

**Supplementary Table 2:** The six potential Batten disease protein biomarkers identified by our pig plasma model were cross-referenced with previously reported plasma proteome data obtained through standard plasma proteomics workflows^15^These workflows encompassed various approaches: “Neat” denotes a neat plasma digestion workflow, “Depleted” signifies the use of a plasma depletion strategy, “Deep Fractionation” involves a high-pH fractionation of depleted plasma, achieved by concatenating 19 fractions into 9, “Proteograph” represents a comprehensive five-NP workflow. In all these workflows, Data-Independent Acquisition (DIA) was utilized to analyze a pooled plasma sample. Notably, our pig plasma model demonstrated its capability to quantify novel putative disease biomarkers that are often undetectable using standard plasma proteomics workflows.

|  | Neat | Depleted | Deep<br>Fractionation | Proteograph | Pig plasma |
| --- | --- | --- | --- | --- | --- |
| **CTSB** |  | X | X | X | X |
| **MHY7** |  |  |  | X | X |
| **CTSS** |  | X | X | X | X |
| **NFL** |  |  |  |  |  |
| **MYL2** |  |  |  |  | X |
| **IFI30** |  |  |  |  | X |

**Supplementary Table 3:** Cellular components associated with genotype at 6-, 24-, and 36-months using the DAVID functional annotation tool.

| DAVID - Cellular Component |  |  |  |  |  |
| --- | --- | --- | --- | --- | --- |
| Age | Related Terms | Count | % | P-value | Benjamini |
| **6 Months** | Cytoplasm | 36 | 20.6 | 4.8E-9 | 7.3E-8 |
|  | Proteasome | 9 | 5.1 | 5.7E-9 | 7.3E-8 |
|  | Secreted | 26 | 14.9 | 7.6E-8 | 6.6E-7 |
|  | Nucleosome core | 4 | 2.3 | 8.1E-3 | 5.3E-2 |
|  | Nucleus | 24 | 13.7 | 1.9E-2 | 9.7E-2 |
|  | Thick filament | 2 | 1.1 | 3.9E-2 | 1.7E-1 |
|  | Mitochondrion | 8 | 4.6 | 5.3E-2 | 2.0E-1 |
|  | Endoplasmic reticulum | 7 | 4.0 | 9.6E-2 | 3.1E-1 |
| **24 Months** | Secreted | 42 | 22.5 | 6.0E-19 | 1.2E-17 |
|  | Proteasome | 9 | 4.8 | 1.1E-8 | 1.1E-7 |
|  | Cytoplasm | 27 | 14.4 | 7.7E-4 | 5.1E-3 |
|  | Lysosome | 4 | 2.1 | 7.5E-2 | 3.7E-1 |
| **36 Months** | Proteasome | 12 | 10.4 | 4.0E-15 | 8.5E-14 |
|  | Secreted | 23 | 20.0 | 9.8E-9 | 1.0E-7 |
|  | Lysosome | 7 | 6.1 | 4.5E-5 | 3.2E-4 |
|  | Cytoplasm | 21 | 18.3 | 4.1E-4 | 2.1E-3 |
|  | Mitochondrion outer membrane | 3 | 2.6 | 4.7E-2 | 2.0E-1 |
|  | Cytoplasmic vesicle | 4 | 3.5 | 7.6E-2 | 2.7E-1 |

**Supplementary Table 4:** Biological processes associated with genotype at 6-, 24-, and 36-month using the DAVID functional annotation tool.

| DAVID - Biological Process |  |  |  |  |  |
| --- | --- | --- | --- | --- | --- |
| Age | Related Terms | Count | % | P-value | Benjamini |
| **6 Months** | Tricarboxylic acid cycle | 4 | 2.3 | 6.5E-4 | 3.1E-2 |
|  | Protein biosynthesis | 6 | 3.4 | 2.2E-3 | 5.3E-2 |
|  | Isoprene biosynthesis | 2 | 1.1 | 5.7E-2 | 6.8E-1 |
|  | Chemotaxis | 3 | 1.7 | 5.8E-2 | 6.8E-1 |
| **24 Months** | Blood coagulation | 4 | 2.1 | 6.0E-4 | 1.6E-2 |
|  | Hemostasis | 4 | 2.1 | 6.9E-4 | 1.6E-2 |
|  | Immunity | 9 | 4.8 | 3.8E-3 | 6.0E-2 |
|  | Chemotaxis | 4 | 2.1 | 7.2E-3 | 8.5E-2 |
|  | Host-virus interaction | 2 | 1.1 | 9.8E-2 | 9.2E-1 |
| **36 Months** | Immunity | 12 | 10.4 | 2.6E-6 | 8.0E-5 |
|  | Innate immunity | 9 | 7.8 | 2.9E-5 | 4.4E-4 |
|  | Chemotaxis | 5 | 4.3 | 2.4E-4 | 2.4E-3 |
|  | Sphingolipid metabolism | 2 | 1.7 | 8.9E-2 | 6.9E-1 |

## Supplementary Info

Formula for calculating sPLS score from normalized data: First protein component: *t*^(*P*)^= 0.983*CTSS + 0.236*CTSB - 0.096 First metabolite component: *t*^(*M*)^ = 0.355*GPE + 0.864*GPI – 0.121 sPLS score = *t*^(*M*)^ + *t*^(*P*)^ = 0.355*GPE + 0.864*GPI + 0.983*CTSS + 0.236*CTSB – 0.217

## Notes

### Competing Interest Statement

The authors have declared no competing interest.

